# A new twist on bacterial motility – two distinct type IV pili revealed by cryoEM

**DOI:** 10.1101/720938

**Authors:** Alexander Neuhaus, Muniyandi Selvaraj, Ralf Salzer, Julian D. Langer, Kerstin Kruse, Kelly Sanders, Bertram Daum, Beate Averhoff, Vicki A. M. Gold

## Abstract

Many bacteria express flexible protein filaments on their surface that enable a variety of important cellular functions. Type IV pili are examples of such filaments and are comprised of a helical assembly of repeating pilin subunits. Type IV pili are involved in motility (twitching), surface adhesion, biofilm formation and DNA uptake (natural transformation). They are therefore powerful structures that enable bacterial proliferation and genetic adaptation, potentially leading to the development of pathogenicity and antibiotic resistance. They are also targets for drug development.

By a complement of experimental approaches, we show that the bacterium *Thermus thermophilus* produces two different forms of type IV pilus. We have determined the structures of both and built atomic models. The structures answer key unresolved questions regarding the molecular architecture of type IV pili and identify a new type of pilin. We also delineate the roles of the two filaments in promoting twitching and natural transformation.

## Main

Type IV pili (T4P) are flexible extracellular protein filaments found on many bacteria. They form multifunctional fibres involved in twitching motility, adhesion, immune evasion, bacteriophage infection, virulence and colony formation. T4P have also been linked to DNA uptake, called natural transformation, which is a powerful mechanism that enables genetic adaptation ^1–3^. The filaments are homopolymers composed of thousands of pilin subunits, which form helical arrays measuring several micrometres in length. How T4P are involved in seemingly unrelated functions such as motility and natural transformation is so far unclear.

Depending on the bacterial species, pilins range from 90 to 250 amino acids in length. They are produced as prepilins with a typical class III signal peptide ^4,5^. The preprotein is translocated via the Sec pathway into the cell membrane where the signal peptide is cleaved by prepilin peptidase, priming the pilin for incorporation into the growing pilus. Filament assembly is ATP-dependent and occurs at an inner membrane platform which contains PilC, PilM, PilN and PilO ^6^. In *Thermus thermophilus*, assembly of pilins into a T4P filament depends on the assembly ATPase PilF, which interacts with the inner membrane platform via PilM ^7^. Two retraction ATPases, PilT1 and PilT2, are essential for T4P depolymerisation ^8,9^. T4P are extruded by the outer membrane secretin PilQ ^10–13^. Recently, it has been suggested that expression of the *T. thermophilus* major pilin PilA4 is temperature dependent, leading to hyperpiliation at suboptimal growth temperatures ^14^. The first two *in situ* structures of T4P assembly machineries were solved only recently in both open (pilus assembled) and closed (pilus retracted) states ^11,15^, yet detailed information regarding the molecular interactions governing filament assembly was lacking.

Crystal structures of full length pilins or head domains from various bacteria are available in different oligomeric states ^6,16–22^. Pilins have a conserved N-terminal α-helix, with a 4-5 stranded anti-parallel β-sheet at the C-terminus. The α-helix forms the core of the filament, while the globular β-sheet head domain is solvent exposed and subject to post-translational modifications ^16,17^. To date, five low-resolution electron cryo-microscopy (cryoEM) structures of isolated T4P have been reported. The first, a 12.5 Å structure from *Neisseria gonorrhoeae*, was sufficient to place crystal structures of pilins into the data but not to resolve their structure within the map ^17^. Four subsequent structures from *Pseudomonas aeruginosa*, two *Neisseria* species and enterohemorrhagic *Escherichia coli* have been determined in the 5-8 Å resolution range ^23–25^.

In this study, we combine different modes of cryoEM: electron cryo-tomography (cryoET) and single-particle cryoEM, with functional data to study the T4P of *T. thermophilus*, which is a well-established model organism. Surprisingly, we detect two forms of T4P, a wider and a narrower form, assembled through the same machinery. We determine structures of the two filaments at so-far unprecedented resolution (3.2 Å and 3.5 Å, respectively). This has enabled us to visualise near atomic-level detail and build atomic models for each filament *ab initio*. Our data unambiguously demonstrate that the wider pilus is composed of the major pilin PilA4. Proteomics and knock-out mutants reveal that the narrow pilus consists of a previously unknown pilin, which we name PilA5. Functional experiments confirm that PilA4 is involved in natural transformation whereas PilA5 is essential for twitching motility ^26^. Our results shed new light on bacterial motility and gene transfer, and will help to guide the development of new drugs to fight microbial pathogens.

### Two types of T4P are assembled from the same machinery

Cells of *T. thermophilus* strain HB27 assemble T4P pili on their surface ^27^, predominantly at the cell poles ^11^. Performing cryoET on cells grown at the optimal growth temperature of 68 °C revealed two types of pilus, with differences in their diameter. They emerge from the same assembly machinery (Fig. 1a, b), suggesting that they are both T4P. We have previously shown that transcription of the major pilin gene, *pilA4*, is upregulated at low temperature ^14^. To address the question of whether the growth temperature affects the assembly of the two forms of pilus, we analysed the pili emerging from cells grown at the sub-optimal growth temperature of 58 °C by cryoET. Again, two types of pilus were observed (Fig. 1c, d). Pili emerge from T4P complexes only sporadically ^11^, thus filaments were isolated from cells in order to investigate their structure in more detail. Both wide and narrow forms of the filament were detected in these preparations (Fig. S1a-d).

**Fig. 1:**
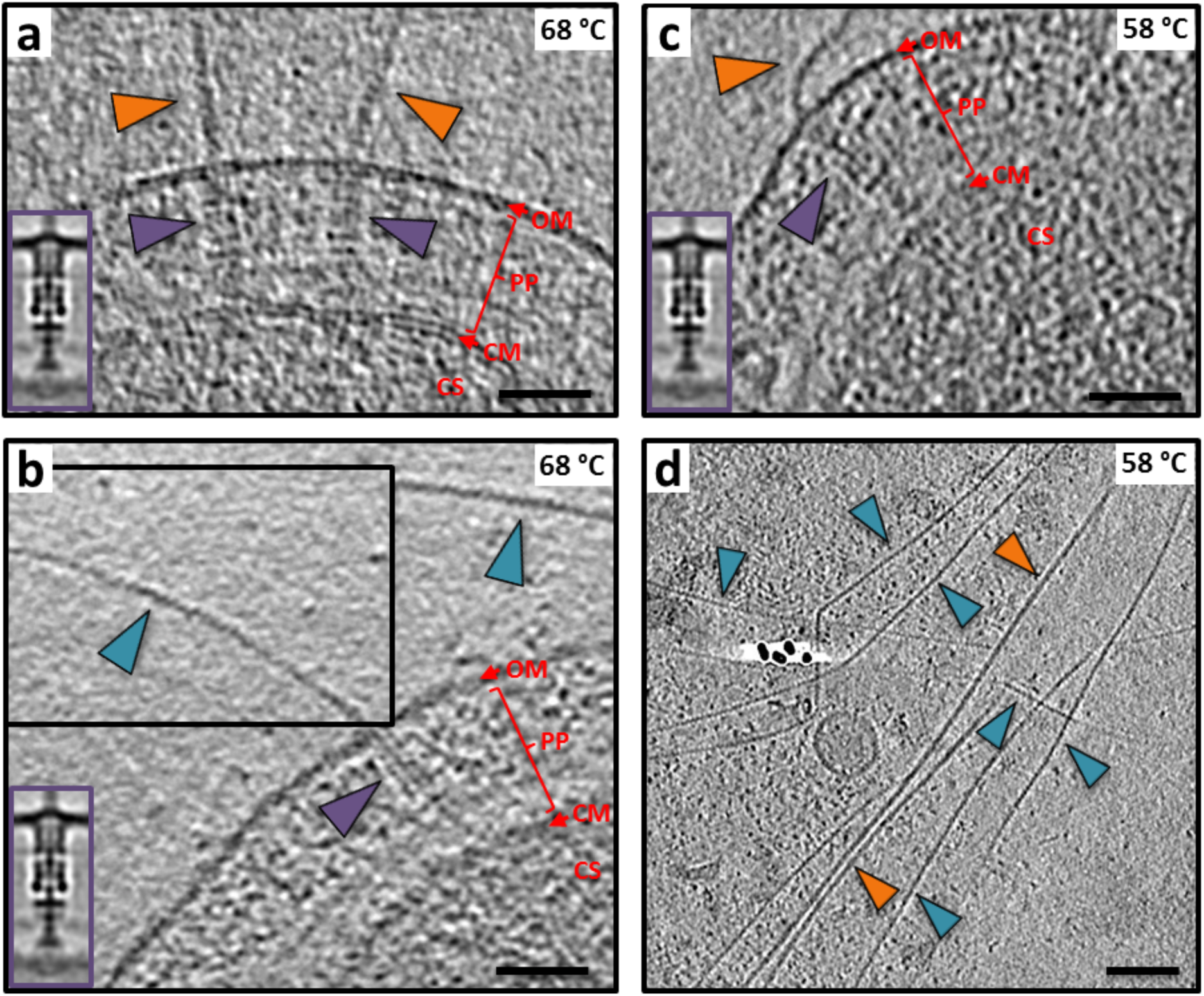
The T4P machinery assembles two types of pilus. **a, b,** Tomographic slices through *T. thermophilus* cells grown at 68 °C show both wide (orange arrowheads) and narrow (teal arrowheads) pili emerging from the T4P machinery (purple arrowheads). The pilus emerges from the cell at an acute angle in (**b**), thus the tomographic volume has been rotated to align with the T4P machinery for visualisation (upper inset box). **c,** Tomographic slice through a *T. thermophilus* cell grown at 58 °C shows a wide pilus (orange arrowhead) emerging from the T4P machinery (purple arrowhead). **d,** Tomographic slice of an area containing many pili from a cell preparation grown at 58 °C. Both wide (orange arrowheads) and narrow pili (teal arrowheads) are visible. Purple insets in **a-c** show the subtomogram average of the T4P machinery (EMD-3023) ^11^. OM, outer membrane; PP, periplasm; CM, cytoplasmic membrane; CS, Cytosol. Scale bars, 50 nm.

To investigate the composition of the two pilus forms, we performed a quantitative bottom-up proteomics analysis. Protein abundance was evaluated by Label-Free Quantitation (LFQ) to determine relative enrichment or loss of particular pilus proteins at either 68 °C or 58 °C. At both temperatures, the major pilin subunit PilA4 was identified as the most abundant protein component (Fig. S2a). The amount of PilA4 increased significantly at 58 °C, in line with the hyperpiliation phenotype ^14^. The second most abundant protein was the uncharacterised protein TT_C1836; the relative abundance was also significantly increased at 58 °C.

To refine the identification of pilins, we performed gel based proteomic analysis (Fig. S2b, Table S1). In order to increase the hyperpiliation phenotype, we further reduced the growth temperature to 55 °C. At this temperature we could identify PilA4 and TT_C1836 as the most abundant proteins in the lower molecular weight bands, likely representing the pilin monomers. In contrast, at the optimal growth temperature of 68 °C, only PilA4 was identified reliably. We questioned whether the two T4P were expressed due to differences in temperature or growth phase. To quantify the abundance of different filaments, cells were grown under different conditions and analysed in the electron microscope. Pili at both cell poles were selected for 2D classification, which enables grouping of similar structures (Fig. S3a). For cells grown on plates, both wide and narrow pili were present at a similar level, whereas the ratio of the two was shifted towards the wider form for cells grown in liquid medium. At 55 °C, only the total number of pili per cell increased, while the ratio between wide and narrow pili was similar between the two temperatures (Fig. S3b).

To analyse the role of PilA4 and TT_C1836 in pilus assembly, we investigated the number of wide and narrow pili per cell in deletion strains grown in liquid media to exponential phase (Fig. S3c). PilA4 deficient cells (*pilA4∷km*) were not able to assemble any pili reliably, whereas TT_C1836 deficient cells (*TT_C1836∷km*) were only defective in their ability to assemble narrow pili. These findings suggest that PilA4 has a role in producing both pilus forms, while TT_C1836 appears to be crucial for the formation of narrow pili only. However, these data do not allow us to discriminate whether the proteins have structural roles in comprising the filaments, or have a more functional role in their assembly mechanism.

### High-resolution maps determine the 3D structure of both filaments

In order to investigate the architecture and protein composition of T4P at high resolution, both filaments were subjected to analysis by single particle cryoEM and helical reconstruction. In our micrographs, the wide pili appeared very straight while the narrow pili showed a much higher degree of curvature, up to 2 μm radius (arrowhead in Fig. S4A, B). Based on 2D classes (Fig. S4c, d) we determined the helical parameters for the wide and the narrow pilus (Fig. S5a– d, see methods section for details). The rise and twist of the wide pilus measured 9.33 Å and 92.5° respectively, and the narrow pilus had a rise of 11.30 Å and a twist of 84.3°. Helical reconstruction ^28^ resulted in maps at 3.22 Å (wide) and 3.49 Å (narrow) (Fig. 2a, b, S5e, f). The diameter of the wide fibre is 70 Å (Fig. 2a) and is roughly cylindrical. In contrast, the narrow filament has a zigzag-like appearance in projection, owing to a 15 Å-wide groove that winds through the fibre. The diameter at any position along the long axis of the fibre axis is therefore only 45 Å (Fig. 2b). Both structures were in good agreement with the data obtained *in situ* by cryoET (Fig. 1). A low-resolution structure of the T4P from *T. thermophilus* was previously determined by cryoET and sub-tomogram averaging, with a diameter of ~3.5 nm ^11^. It seems likely that this conformation represents the narrower form of the pilus.

**Fig. 2:**
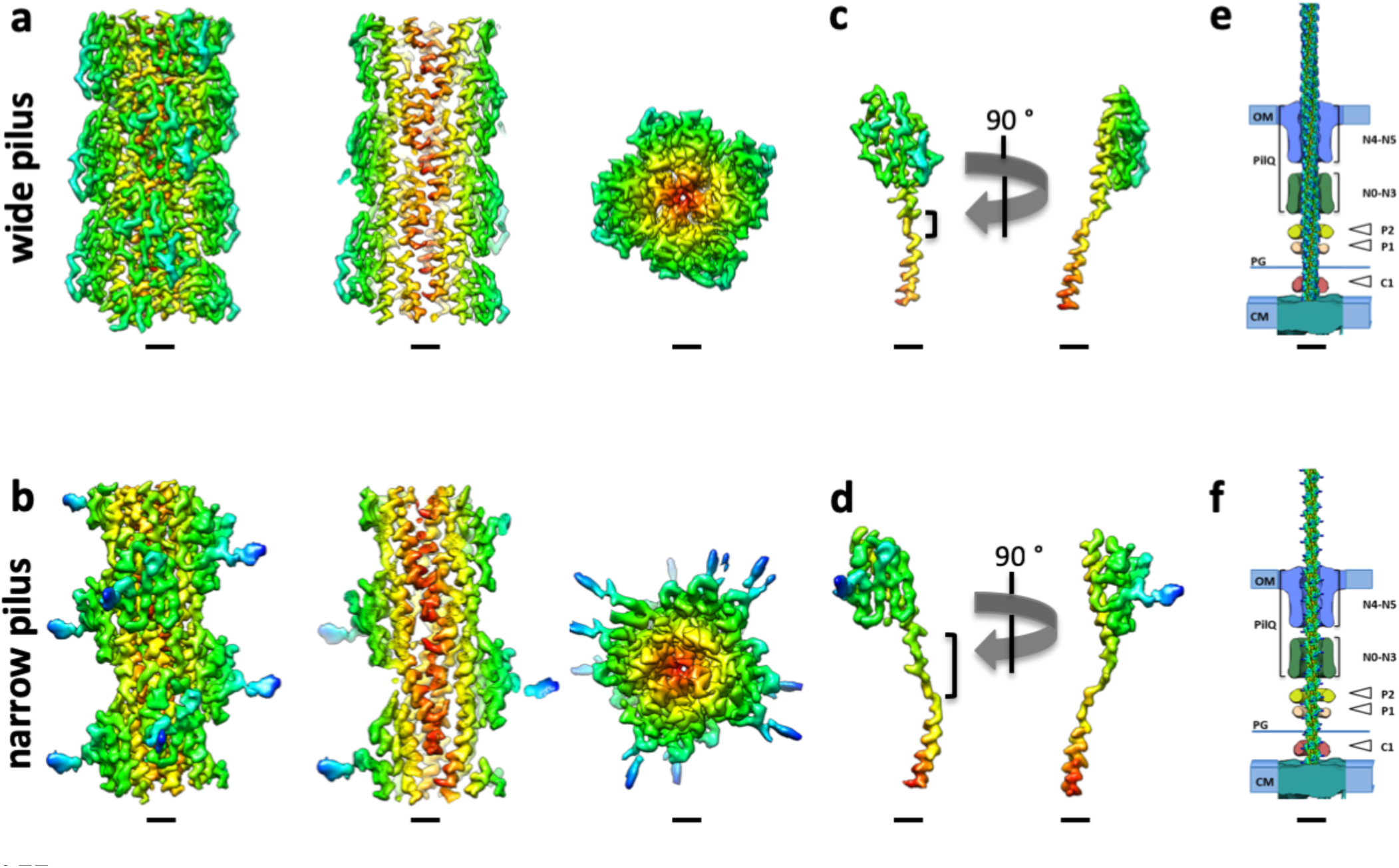
Helical reconstruction of *Tth*HB27 pili. **a, b,** Side views, cross sections and top views of the density maps of the wide (**a**) and narrow (**b**) pilus fibres. Coloured by cylinder radius from red (centre) to blue (periphery). Scale bars, 10 Å. **c, d,** A single subunit of the wide (**c**) and narrow (**d**) pilus segmented from the maps in **a** and **b**. The brackets show the melted stretch in the N-terminal α-helix. Scale bars, 10 Å. **e, f,** Pilus filaments docked into the subtomogram average of the open state of the T4P machinery (EMD-3023) ^11^. Domains of the secretin PilQ (blue and green; N0-N5), with unassigned protein densities P2 (yellow), P1 (orange) and C1 (red) are shown. OM, outer membrane; PG, peptidoglycan; CM, cytoplasmic membrane. Scale bars, 10 nm.

Both maps clearly resolved individual pilin monomers. The peptide backbone could be easily traced throughout each subunit and large side chains were visible (Fig. 2c, d). The centre of each filament is formed by a bundle of long N-terminal α-helices, as has been demonstrated for other filaments ^17,23–25,29^. In both maps, each α-helix is interrupted by an unfolded stretch (brackets in Fig. 2c, d), a conserved feature observed in the N-terminal domains of all available T4P structures. Interestingly, the unfolded region is significantly longer for the pilin comprising the narrow filament, resulting in a longer N-terminal stalk. The outer regions of both filaments are formed by globular domains consisting of β-strands, a typical hallmark of the T4P C-terminal domain ^30^. Whilst the C-terminal head domains of both pili are comprised of central β-sheets, the domain size and the region linking to the N-terminal α-helix appear different. These findings suggest that the two pili are not only distinct with regards to their helical parameters, but also consist of different proteins.

As shown, we observe both filaments emerging from the T4P machinery (Fig. 1). This is supported by previous studies showing that mutants defective in the PilQ secretin channel do not extrude pili ^27^. In addition, previous studies have shown that a mutant defective in the assembly ATPase PilF is non-piliated ^9^. This suggests that both pili are extruded by the same assembly machinery. The aperture within the central channel of PilQ is of sufficient dimension to accommodate either form (Fig. 2e, f).

### Atomic models of T4P

The resolution and quality of both maps allowed us to unambiguously build an atomic model for each filament *ab initio* (Fig. 3a - f). Guided by our mass spectrometry results, the position of large side chains and clear differences in the length of the polypeptide backbones, we were able to identify PilA4 as the building block for the wide pilus and the previously uncharacterised protein TT_C1836 as the subunit for the narrow filament. We now propose that TT_C1836 be named PilA5, in keeping with *Thermus* nomenclature.

**Fig. 3:**
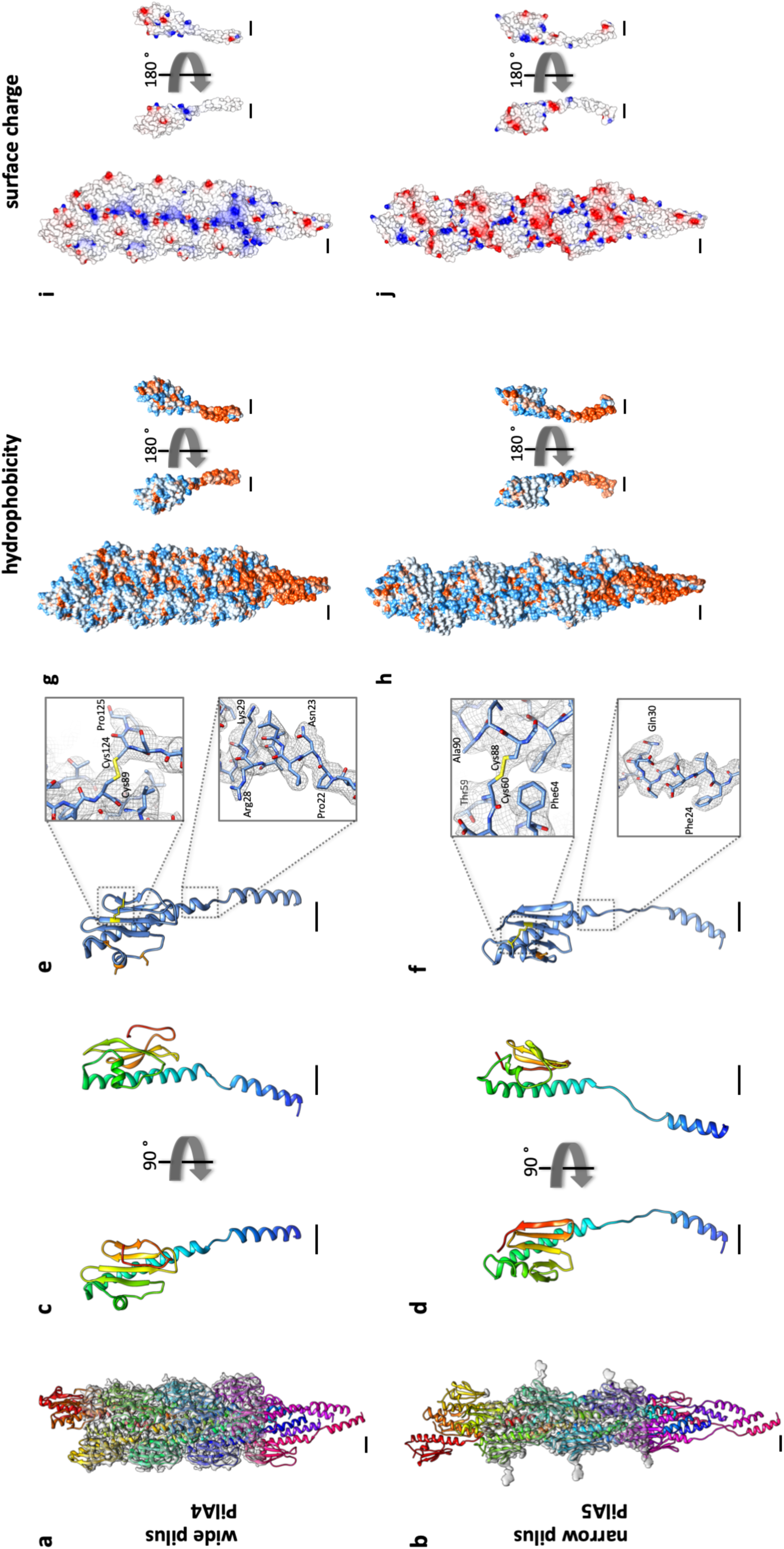
Molecular models of wide and narrow pili. **a, b,** Molecular models of short sections of filaments (15 subunits each) with the corresponding EM density maps. **a,** wide pilus comprised of PilA4; **b,** narrow pilus comprised of PilA5. Scale bars, 10 Å. **c, d,** Ribbon representation of a single PilA4 (**c**) and PilA5 (**d**) subunit. N-terminus blue, C-terminus red. Scale bars, 10 Å. **e, f,** Ribbon representation of a single PilA4 (**e**) and PilA5 (**f**) subunit with selected sidechains shown. Yellow, disulphide bond; orange, serines with posttranslational modification. Insets show details of the molecular model within the EM density map. Top insets, disulphide bond; bottom insets, side chain densities. Scale bars, 10 Å. **g, h,** Hydrophobicity of PilA4 (**g**) and PilA5 (**h**) filaments (left) and single subunits (right). Hydrophobic (red) and hydrophilic (blue) residues are shown. Scale bars, 10 Å. **i, j,** Electrostatic surface charge of PilA4 (**i**) and PilA5 (**j**) filaments (left) and single subunits (right). Negative charges (red) and positive charges (blue) are shown. Scale bars, 10 Å.

The N-terminal α-helix, including the unfolded stretch, is comprised of the first 54 (PilA4) or 53 amino acids (PilA5). In both proteins the helix is disrupted by an unfolded stretch around the conserved Pro22 (Fig. 3c, d, S6). The stretch in PilA4 is 4 amino acids long as opposed to 10 amino acids long in PilA5. The region between the N-terminal α-helix and the C-terminal β-sheet, the so-called glycosylation loop, ranges in PilA4 from amino acids 55 to 77, with a two-turn α-helix comprising amino acids 61-67. The C-terminal region is an antiparallel four-stranded β-sheet with the last strand facing towards the N-terminus followed by a loop that ends on the β-sheet. A disulphide bond between Cys89, which is located in the second strand, and the penultimate amino acid Cys124, likely stabilises the C-terminus (Fig. 3e). The glycosylation loop in PilA5 spans amino acids 54 to 71, with amino acids 62 to 65 forming a one-turn helix. The C-terminal β-sheet is composed of five strands, one more than observed in PilA4. Due to the additional β-strand in PilA5, the last strand faces away from the N-terminus of the protein. The C-terminus of PilA5 is located between the β-sheet, the glycosylation loop and the long α-helix. A disulphide bond is formed between Cys60 in the glycosylation loop and Cys88 in the third β-strand (Fig. 3f). Both pilins are highly hydrophobic at the N-terminal part of the α-helix, and more hydrophilic on the surface of the globular domain (Fig. 3g & h). The hydrophobic helices bundle and form the hydrophobic core of the assembled filament. PilA4 has no net charge but the filament displays a distinctive positively charged groove along the filament axis (Fig. 3i). In contrast, PilA5 has a total of 2 negative charges per subunit which leads to a patch of negative charge winding around the filament (Fig. 3j).

A network of cooperative interactions between pilin subunits holds the fibres together. Each subunit has 6 (wide form) or 7 (narrow form) physical interaction partners in each direction of the fibre (thus 12 or 14 total interaction partners) spread in side-by-side or top-to-bottom directions (Fig. S7a, b). Most of the interactions involve a large portion of the N-terminal α-helices within the hydrophobic core as well as the head domains. In the wide (PilA4) filament, each subunit (subunit A) interacts with the N-termini that project down from the next two subunits (B and C) above and from the subunits which are six and seven subunits above (G and H). A second interaction takes place between the upper part of the α-helix in subunit A and the glycosylation loop in subunit B (Fig. S7a). In the narrow (PilA5) filaments, subunit A interacts via its α-helix with the N-termini of subunits B, C, F, G and H, while there is an additional interaction between the upper part of the α-helix in subunit A with the glycosylation loop in subunit B (Fig. S7b).

For both types of pili, the largest interaction interface is between subunits which are 3 or 4 subunits apart (subunit A with subunit D and E). Thus each pilin subunit has a large interaction interface with 6 other subunits (B, D and E in Fig. S7a, b) and 6 or 8 smaller interaction sites (C, G, H in Fig. S7a and C, G, H, F in Fig. S7b). Most interactions involve the hydrophobic sidechains in the centre of the filament and appear to be nonspecific, likely allowing sliding movements between the subunits when the filaments are stretched. This is in accordance with the observation that pili can stretch up to threefold upon force ^31^. In addition, PilA4 contains salt bridges between Asp53 in subunit A and Arg30 in subunit D, and between Glu48 in subunit A and Arg28 in subunit E. In contrast, PilA5 contains a single salt bridge between Glu68 in subunit A and Arg23 in subunit D (Fig. S7c). The conserved Glu5 is likely required to neutralise the positive charge of the N-terminus within the hydrophobic core of the filament ^4,32,33^. A salt bridge is also found between Glu5 and the N-terminus of the neighbouring subunit in other T4P ^23,24^. For both *T. thermophilus* fibres the distance between Glu5 and adjacent N-termini is too far to form a salt bridge. Instead, Glu5 forms an intramolecular salt bridge to the N-terminus in the same subunit (Fig. S7c). This was also modelled for the related *Klebsiella oxytoca* pseudopilus ^34^.

### Posttranslational modification

Densities were observed in both EM maps that protrude into the solvent and cannot be attributed to the polypeptide backbone and were too large to account for an amino acid side chain (Fig. 4). Interestingly, these densities co-localised with serine residues and were similar in appearance to previously published densities attributed to glycosylation sites ^29,35,36^. We suggest that these densities correspond to O-linked glycosylation, which is consistent with the previous finding that the major pilin PilA4 of *T. thermophilus* is glycosylated ^37^. Glycosylation has also been observed in similar locations in the X-ray structure of *N. gonorrhoeae* type IV pilin head domain (PDB: 2HI2) ^16,17,38^. In PilA4, we found extra densities at 3 serine residues in the glycosylation loop (Ser59, Ser66 and Ser71), while only one serine appeared to be modified in PilA5 (Ser73). Interestingly the density was much more pronounced in PilA5 than in PilA4. This may be due to a less flexible glycan moiety in PilA5, allowing for improved resolution, or by a different composition of the sugar residues entirely. Glycosylation may enhance temperature stability via additional hydrogen bonds, increase adhesive properties (either to surfaces or to small molecules such as DNA) or act as recognition tags for cell-cell communication ^39^.

**Fig. 4:**
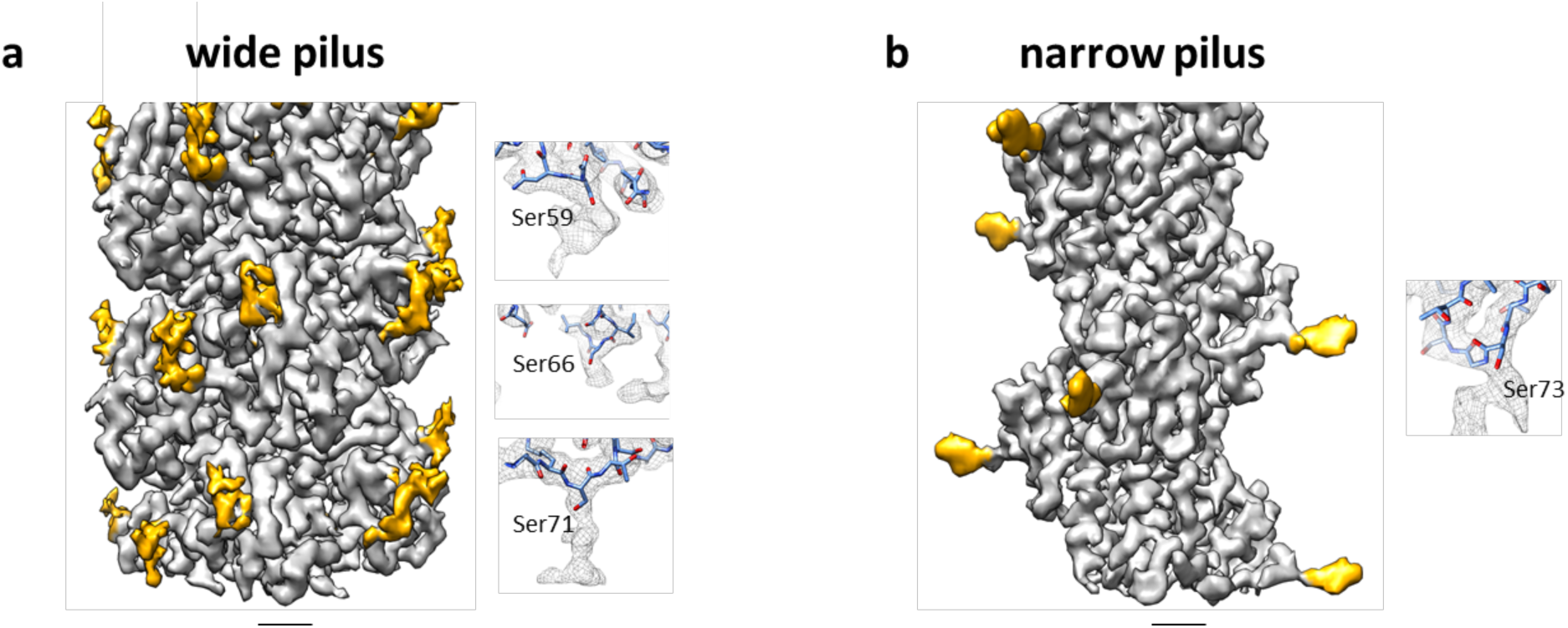
Post-translational modification. **a, b,** Surface representation of the EM maps of the wide (**a**) and narrow (**b**) pilus showing the protein model (grey) and densities that protrude into the solvent and could not be attributed to the polypeptide backbone or an amino acid side chain. These are most likely glycans (yellow). Insets show close-ups of large unassigned densities near serine residues. Scale bars, 10 Å.

### The functional importance of two types of pili

A key outstanding question pertains to the functional relevance of the two types of T4P. To investigate this question, we performed various functional analyses on PilA4 and PilA5 deletion strains. We assessed cell lines without pili (*pilA4∷km*), with wide pili only (*pilA5∷km*), or a mixed population of wide and narrow pili (wild type).

We analysed cellular motility by twitching assays at 68 °C and 55 °C. Wild-type cells formed characteristic twitching zones of ~2 cm and ~1.2 cm in diameter, respectively. The mutants *pilA4∷km* and *pilA5∷km* did not exhibit any twitching motility (Fig. 5a). Since the immotile *pilA5*∷km cells could still produce wide pili comprised of PilA4, we deduce that PilA5 is required to promote cell movement. Cells lacking both types of pili in the *pilA4∷km* mutant were completely defective in natural transformation, in agreement with our former finding ^40^. Transformation efficiency was only partially reduced in the *pilA5∷km* mutant (~30%), which expresses wide PilA4 pili (Fig. 5b). This corresponds to our previous finding that a *pilA5∷km* mutant is still transformable ^40^ and demonstrate that the narrow pili are dispensable for DNA uptake.

**Fig. 5:**
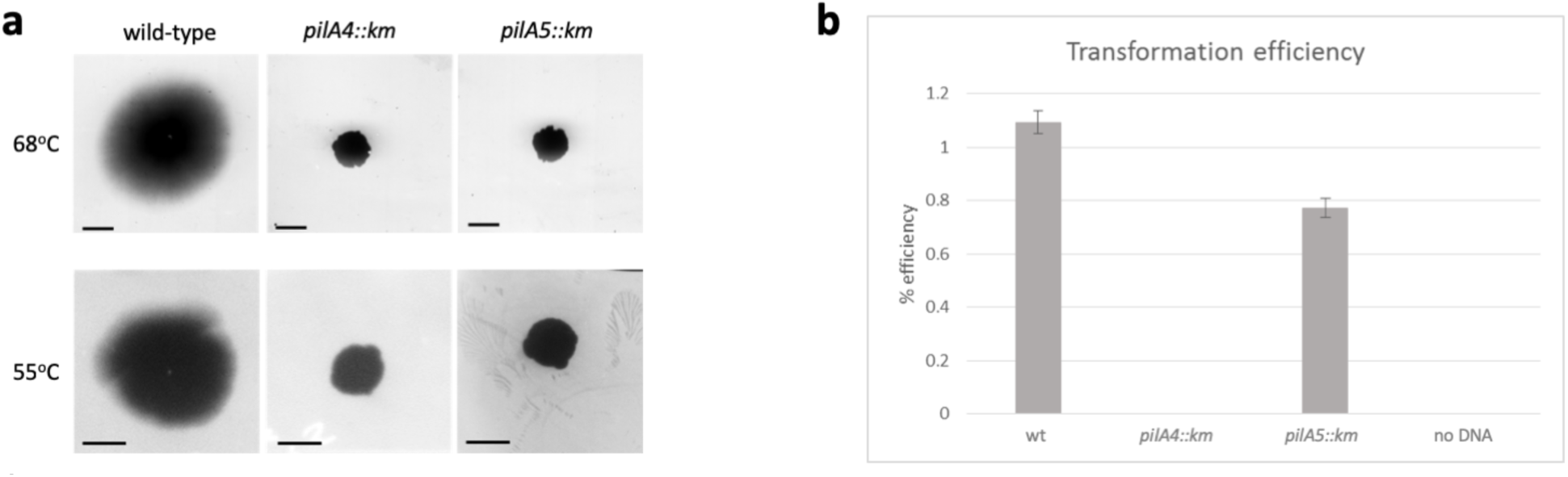
Functional characterisation of *T. thermophilus* mutants. **a,** Twitching motility of *T. thermophilus* HB27 strains. Only wild-type cells show twitching. Scale bars = 0.5 cm. **b,** Natural transformation efficiency of *T. thermophilus* HB27 strains.

In summary, we conclude that PilA4 has three known roles: it promotes pilus formation for both wide and narrow filaments, it comprises the main structural element of the wide T4P, and it plays a role in natural transformation. We determine two functions for PilA5: it forms the basis of the narrow T4P and is a requirement for cell motility.

## Discussion

We have determined the first cryoEM structures of T4P that have allowed atomic models to be built *ab initio*. Moreover, we have discovered two distinct T4P filaments, which are composed of different proteins. Our data provide compelling evidence that PilA5 is essential for twitching motility and confirm the previous finding that PilA4 is involved in natural transformation ^40^. In addition, we find that PilA4 is essential for the assembly of both wide and narrow pili. PilA4 may therefore play a crucial regulatory role, could initiate pilus formation, or even form a capping structure. In many bacteria, minor pilins are thought to prime pilus assembly by reducing the energy barrier to the extraction of pilins from the membrane ^41^. In *Thermus*, PilA4 may perform this role.

The unique functionality of PilA4 and PilA5 is hardcoded in their distinct structural features. Both filaments follow the conserved T4P blueprint, encompassing a central bundle of hydrophobic N-terminal α-helices and a hydrophilic C-terminal β-strand globular domain ^39^. Structural variations in PilA4 and PilA5 determine distinct inter-subunit interactions, helical parameters, mechanical properties, adhesiveness and binding affinity. Narrow pili comprised of PilA5 are more flexible than those comprised of PilA4. In line with their predicted role in twitching, narrow pili that can bend and flex would enable the filaments to curve from the surface of cells to interact with surfaces, negotiate obstacles and increase the exploratory range of the cell. Their overall net negative surface charge would enhance the adhesive properties of the fibre and facilitate surface adhesion.

In accordance with the role of PilA4 in natural transformation, the surface of wide PilA4 filaments show a striking line of positively-charged residues along the long axis. We speculate that these may be involved in binding the negatively charged DNA backbone. A double stranded DNA molecule fits into a right-handed helical groove flanked by glycan residues. This would allow interaction of the DNA backbone phosphates with positive charges in the PilA4 filament at regular intervals. Flanking glycans and negative charges may coordinate proper DNA binding (Fig. 6). This suggestion is in good agreement with the well characterised roles of T4P in DNA binding and uptake in *Neisseria*, and of DNA binding in *P. aeruginosa* ^42,43^. Our proposal is also consistent with that of Craig *et al*, who suggest that the positively charged groove of gonoccocal T4P is wide enough to bind the negatively charged backbone of dsDNA^17^. Interestingly, previous experiments have shown that pili are not solely important for natural transformation. For example, a mutant defective in the PilF assembly ATPase was impaired in piliation and was hypertransformable, and a mutant carrying a deletion in a domain of the secretin PilQ was impaired in piliation but exhibited wildtype transformation frequencies ^10,44^. Taken together, we suggest that wide pili comprised of PilA4 may capture DNA rather like a fishing net, thus improving the efficiency of DNA uptake by increasing the local concentration of DNA near the outer membrane.

**Fig. 6:**
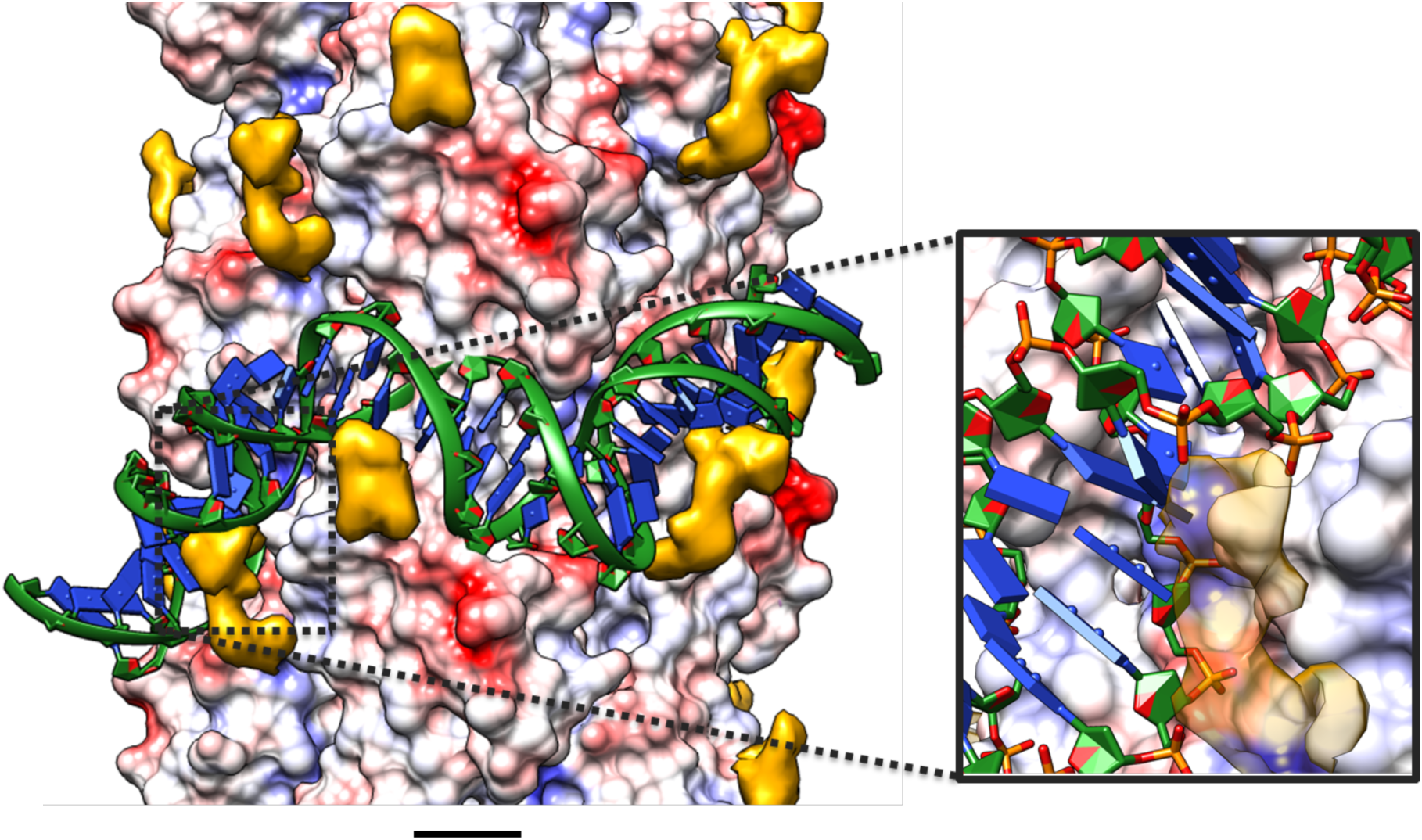
Proposed DNA binding to wide pili. A double stranded DNA molecule (green) is modelled around a wide pilus shown in surface charge representation (negative charges, red; positive charges, blue). The DNA backbone fits neatly into the positively charged groove of the PilA4 filament (inset). Post-translational modifications are shown in yellow (transparent yellow in inset). Scale bar, 10 Å.

In evolutionary terms, bacteria appear to have reused the T4P blueprint to develop a system that can assemble two different filaments with unique properties. This could enable tasks to be performed more effectively and at reduced energy cost to the cell. It will now be interesting to discover if this principle occurs in other bacterial species, and excitingly, will open avenues to the development of vaccines or therapeutics targeting a particular T4P mechanism.

## Experimental procedures

### Cultivation of organisms

*T. thermophilus* HB27 was grown in TM^+^ medium (8 g/l tryptone, 4 g/l yeast extract, 3 g/l NaCl, 0.6 mM MgCl_2_, 0.17 mM CaCl_2_) ^45^ at 55 °C, 58 °C or 68 °C. Antibiotics were added when appropriate (kanamycin, 80 mg/ml; streptomycin, 100 mg/ml in solid medium (containing 2 % agar [w/vol]) or kanamycin, 60 mg/ml; streptomycin 100 mg/ml in liquid medium). Disruption of *TT_C1836* and *pilA4* was performed by insertion of a kanamycin marker ^27^.

### Purification of pili

*T. thermophilus* HB27 cells were grown for two days on TM^+^ medium at 68 °C or three days at 55 °C.. Cells were scraped off and resuspended by pipetting and shaking in ethanolamine buffer (0.15 M ethanolamine, pH 10.5). Cells were sedimented by centrifugation (16.200 *x g*, 1 h, 4 °C). The supernatant was gently mixed with saturated ammonium sulfate solution [10/1 (v/v)] and incubated on ice for 12 h. Pili were precipitated by centrifugation for 10 min (16,200 *x g*, 4 °C). The resulting pellet was washed twice with TBS buffer (50 mM Tris/HCl, 150 mM NaCl, pH 7.5). The pellet was resuspended by incubation in distilled water for 4 h. 10x buffer (500 mM Tris/HCl, 500 mM NaCl, 10 mM CaCl_2_, 10 mM MgCl_2_, pH 7.5) was added prior to structural analyses.

### CryoET sample preparation and imaging

Cubes of agar with growing *T. thermophilus* HB27 cells were cut out, placed into EDTA buffer (20 mM Tris/HCl, 100 mM EDTA, pH 7.4) and gently agitated for 1 hour at room temperature. Samples were mixed 1:1 with 10 nm protein A-gold (Aurion, Wageningen, The Netherlands) as fiducial markers and glow-discharged R2/2 Cu 300 mesh holey carbon-coated support grids (Quantifoil, Jena, Germany) were dipped into the solution. For analysis of isolated pili by cryoET, preparations were mixed 1:1 with 10 nm protein A-gold fiducial markers and solutions were gently pipetted onto the grids. All grids were blotted using Whatman 41 filter paper for ~4 s in a humidified atmosphere and plunge-frozen in liquid ethane in a home-built device or using a Vitrobot Mark IV (Thermo Fisher, Waltham, USA). Grids were maintained under liquid nitrogen and transferred into the electron microscope at liquid nitrogen temperature.

Tomograms were typically collected from +60° to −60° at tilt steps of 2° and 5 - 7 μm underfocus (whole cells), or at 3 μm underfocus (isolated pili), using either a Tecnai Polara, Titan Krios (Thermo Fisher) or JEM-3200FSC (JEOL, Tokyo, Japan) microscope, all equipped with field emission guns operating at 300 keV. All instruments were fitted with energy filters and K2 Summit direct electron detector cameras (Gatan, Pleasanton, USA). Dose-fractionated data (3-5 frames per projection image) were collected using Digital Micrograph (Gatan). Magnifications varied depending on microscope; pixel sizes were within the range 3.8 – 4.2 Å. The total dose per tomogram was < ~140e^−^/Å^2^. Tomograms were aligned using gold fiducial markers and volumes reconstructed by weighted back-projection using the IMOD software (Boulder Laboratory, Boulder, USA) ^46^. Contrast was enhanced by non-linear anisotropic diffusion (NAD) filtering in IMOD ^47^.

### Calculations of pilus diameter

Slices through tomograms were analysed by drawing a plot profile of grey values in ImageJ ^48^, which could be exported as a function of distance. Statistical analysis of pilus diameters was conducted with a sample size of 8 tomograms containing ~60 isolated pili.

### Negative-stain electron microscopy

Two microliters of purified pili were pipetted onto glow-discharged carbon-coated Cu 400 mesh support grids (Sigma-Aldrich) for 2 minutes. Grids were blotted with Whatman No 41 filter paper and stained with 5 % ammonium molybdate for 60 s. Images were recorded with a Tecnai Spirit microscope (Thermo Fisher) operated at 120 keV and a OneView CMOS camera (Gatan). Images were analysed for fibre quality, size, sample density and homogeneity using EMAN2 ^49^. For whole cell samples of *T. thermophilus*, either liquid culture was used directly or some cells were carefully scraped off the plates and resuspended in TBS. If required, cells were diluted in TBS. Negative staining was performed as described above. For the quantitative analysis of number and type of pili per cell, filaments from each cell pole were counted and helices of equal length were selected using e2helixboxer (EMAN2) and subsequently classified in 2D using RELION ^50^. The percentage of wide, narrow and unassigned pili was calculated based on the number of particles in each class.

### CryoEM sample preparation and imaging of fibres

Three microliters of isolated pilus suspension were pipetted onto a glow-discharged R2/2 Cu 300 mesh holey carbon-coated grids (Quantifoil). Grids were plunge frozen in liquid ethane after blotting using a Vitrobot Mark IV (Thermo Fisher) and stored in liquid nitrogen. Cryo images were collected with a Titan Krios microscope (Thermo Fisher) at the UK national electron bio-imaging centre (eBIC), equipped with a field emission gun operating at 300 keV. The microscope was fitted with K2 Summit direct electron detector and Quantum energy filter (both Gatan, Pleasanton, USA). Dose-fractionated data were collected at 1.5 – 4 μm defocus using EPU (Thermo Fisher). 3138 micrographs containing both forms of pili were collected as 40-frame movies, corresponding to 8 seconds at a frame rate of 1 frame for every 0.2 seconds. The total dose was 48 electrons/Å^2^ at a magnification of 130,000 x, corresponding to a pixel size of 1.048 Å.

### Image processing, symmetry determination and helical reconstruction

Drift correction was performed using UNBLUR ^51^. Straight sections of thin and wide fibres were boxed separately from the drift corrected images using the helixboxer function of EMAN2, such that the filaments were centred in each rectangular box. Helical reconstruction was performed using the boxed filaments and SPRING as follows ^52^. Contrast transfer function (CTF) correction was performed using CTFFIND ^53^. In order to determine the helical parameters of the wide filaments a subset of the boxed filaments were cut into small segments of 373 Å (a multiple of the helical rise) with an 80 % overlap, yielding a total of 9576 segments, which were classed in 2D (Fig. S4a). Close examination of the segments indicated a filament diameter of 75 Å. The calculated power spectrum from the total segments indicated clear layer lines that could be indexed. A meridional reflection at approximately 9 Å and a layer line of order 1 at approximately 36 Å indicated that there are approximately 4 subunits per turn. The ninth layer was found to be of order 1, suggesting that the helix repeats exactly after nine turns, with a non-integer number of subunits per turn. The SEGMENTCLASSRECONSTRUCT module in SPRING on class averages was used to determine the accurate helical symmetry (Fig. S5a). The suggested output was determined to be either 4.10 or 3.89 subunits in a helical pitch of 36.3 Å. In order to determine the helical parameters for the narrow filaments a subset of the boxed filaments were cut into segments of 800 Å with a step size of 330 Å and classed in 2D (Fig. S4b). The helical pitch could be determined directly as 48.1 Å. A meridional reflection and thus a helical rise at 11.3 Å could be identified. These parameters allow calculation of a helical rotation of 84.6° and 4.26 subunits per turn. The SEGMENTCLASSRECONSTRUCT module in SPRING was again used to determine the accurate helical symmetry (Fig. S5b). The suggested output was determined to be either 4.11, 4.14, 4.27 or 4.30 subunits with a helical pitch of 48.1 Å. 3D reconstruction was performed using the above parameters by iterative projection matching and back projection as implemented in the SEGMENTREFINE3D of SPRING, starting from a solid cylinder of 75 Å as a reference. Examination of the Fourier transforms simulated from the reconstructed volume to that experimentally calculated from fibres indicated that 3.89 subunits in a pitch of 36.3 Å (accounting for a helical rise of 9.33 Å and a helical rotation of 92.5 degrees) is correct for the wide filaments, and 4.27 subunits in a pitch of 48.1 Å (accounting for a helical rise of 11.26 Å and a helical rotation of 84.3 degrees) is correct for the narrow filaments (Fig. S5c, d). For the final maps 300 Å segments with a step size of three times the helical rise from 400 images were extracted. Doubling the number of used images did not further increase the final resolution. The calculated final maps were determined at 3.22 Å resolution from 98,415 asymmetric units for wide filaments, and 3.49 Å from 76,866 asymmetric units for narrow filaments (Fig. S5e, f) using Fourier shell correlation (0.143 cut-off). For the final maps a B-factor of −60 Å^2^ was applied. Figures were drawn in Chimera, Coot and CCP4mg ^54–56^.

### Model building

Atomic models for both forms of pili were built manually *de novo* in Coot. We assumed that one of the two pili consists of the major pilin PilA4, which is 125 amino acids in length. The backbone as well as all large side chain densities of PilA4 match the density map of the wider form of the pilus. While tracing the backbone into the density maps it became apparent that the subunits forming the narrower pili are ~10% smaller than the subunits of the wider pili. Following the results of mass spectrometry and deletion experiments we modelled the second most abundant protein, TT_C1836 (111 amino acids) into the density map of thin pili. The backbone and visible side chains fit perfectly into the density map. The structure was iteratively refined by Refmac5 ^57^ followed by manual rebuilding in Coot and ISOLDE ^58^. The final models contain all amino acid residues of the mature protein. The double stranded DNA in Fig. 6 (based on PDB: 1bna) was modelled around the wide pilus using Chimera and Coot.

### Data deposition

The cryo-EM maps were deposited in the Electron Microscopy Data Bank with accession codes EMD-XXXX (wide pilus) and EMD-YYYY (narrow pilus). The structure coordinates of the atomic models of the wide and the narrow pilus were deposited in the Protein Data Bank with accession numbers WWWW and ZZZZ, respectively.

### Sequence Alignment

Sequence alignment was performed using the PRALINE server ^59^ with the default settings (weight matrix: BLOSSUM62, gap opening penalty: 12, gap extension penalty: 1).

### Mass spectrometry

Purified pilus preparations were processed using a modified FASP workflow ^60^ as described previously ^61^. In brief, reduced and alkylated protein extracts were digested sequentially with Lys-C and trypsin on Microcon-10 filters (Merck Millipore, # MRCPRT010 Ultracel YM-10). Digested samples were desalted using ZipTips according to the manufacturer’s instructions, dried in a Speed-Vac and stored at −20 °C until LC/MS-MS analysis. Dried peptides were dissolved in 5 % acetonitrile with 0.1 % formic acid, and subsequently loaded using a nano-HPLC (Dionex U3000 RSLCnano) on reverse-phase columns (trapping column: particle size 3μm, C18, L=20mm; analytical column: particle size <2μm, C18, L=50cm; PepMap, Dionex/Thermo Fisher). Peptides were eluted in gradients of water (buffer A: water with 5 % v/v acetonitrile and 0.1 % formic acid) and acetonitrile (buffer B: 20 % v/v water and 80 % v/v acetonitrile and 0.1 % formic acid). All LC-MS-grade solvents were purchased from Fluka. Gradients were ramped from 4 % to 48 % B in 120 minutes at flow rates of 300 nl/min. Peptides eluting from the column were ionised online using a Thermo nanoFlex ESI-source and analysed in a Thermo “Q Exactive Plus” mass spectrometer. Mass spectra were acquired over the mass range 350-1400m/z (Q Exactive Plus) and sequence information was acquired by computer-controlled, data-dependent automated switching to MS/MS mode using collision energies based on mass and charge state of the candidate ions.

Raw MS data were processed and analysed with MaxQuant ^62^. In brief, spectra were matched to the full 15 nalyse.org database (reviewed and non-reviewed, downloaded on the 13/05/2016) and a contaminant and decoy database. Precursor mass tolerance was set to 4.5 ppm, fragment ion tolerance to 20 ppm, with fixed modification of Cys residues (carboxyamidomethylation +57.021) and variable modifications of Met residues (Ox +15.995), Lys residues (Acetyl +42.011), Asn and Gln residues (Deamidation +0.984) and of N-termini (carbamylation +43.006). Peptide identifications were calculated with FDR = 0.01, and proteins with one peptide per protein included for subsequent analyses. Proteomics data associated with this manuscript have been uploaded to PRIDE ^63^. Anonymous reviewer access is available upon request. Peptide intensities (label free quantitation) were analysed using MaxQuant and Perseus ^62^. Differential abundance of proteins (detected in at least 3 of 4 replicates in each condition) was analysed using a two-sided t test with a FDR of 0.01 and s0= 0.05.

For gel-based MS, purified pili were separated by SDS-PAGE (Mini Protean TGX 4-15 %, Biorad, Hercules, USA) and proteins were stained using Bio-Safe Coomassie Stain (Biorad, Hercules, USA). Bands were cut out and analysed by MS at the University of Bristol Proteomics Facility.

### Twitching motility

*T. thermophilus* HB27 strains were grown at 68 °C for 3 days and at 58 °C for 7 days under humid conditions on minimal medium agar plates ^10^ containing 0.1 % bovine serum albumin. Plates were then stained with Coomassie blue and cells washed off to reveal twitching zones. The contrast has been inverted in the images.

### Natural transformation

*T. thermophilus* wild-type, *pilA4∷km* and *TT_C1836∷km* mutants were cultured in TM^+^ media containing appropriate antibiotics for 24h at 68 °C, 150 rpm (New Brunswick Innova 42, Eppendorf, Hamburg, Germany). These cultures were used to inoculate 10 ml TM^+^ media (with appropriate antibiotics) to a starting OD600 = 0.2 and incubated until OD600 = 0.5 was reached. 30 μl of the cultures were transferred into 370 μl prewarmed TM^+^ medium, and 10 μg genomic DNA from a spontaneous streptomycin-resistant HB27 mutant was added (HB27 Strep). The cultures were incubated for 30 min at 68 °C, 150 rpm. These were subsequently diluted (400 μl in 3 ml TM^+^) and incubated for an additional 3 h at 68 °C, 150rpm. Transformations were plated in suitable dilutions onto TM^+^ agar plates containing 100 μg/ml streptomycin to determine the number of transformants. The number of viable cells was determined by plating on TM^+^ agar plates lacking streptomycin. Following incubation for 2 days at 68°C, colonies were counted. The transformation efficiency was calculated as the number of transformations per number of living cells.

## Acknowledgements

We thank Werner Kühlbrandt and Deryck Mills for their support at the MPI of Biophysics in Frankfurt, and Carsten Sachse for indispensable assistance using SPRING and feedback on early versions of this manuscript. We thank Mathew McLaren for maintenance of the EM facility in Exeter and we acknowledge access and support of the GW4 Facility for High-Resolution Electron Cryo-Microscopy, funded by the Wellcome Trust (202904/Z/16/Z and 206181/Z/17/Z) and BBSRC (BB/R000484/1). We acknowledge Diamond for access and support of the cryoEM facilities at the UK national electron bio-imaging centre (eBIC), proposal EM18258, funded by the Wellcome Trust, MRC and BBSRC. We thank Kate Heesom (University of Bristol Proteomics Facility), Imke Wüllenweber and Fiona Rupprecht (MPI for Biophysics) for MS experiments. We acknowledge the BBSRC (BB/R008639/1), Max-Planck-Society, the University of Exeter and the Deutsche Forschungsgemeinschaft (AV 9/6-2) for funding.

## Author contributions

Major contributions to (i) the conception or design of the study (AN, MS, RS, BD, BA, VAMG) (ii) the acquisition, analysis, or interpretation of the data (AN, MS, RS, KK, KS, JDL, BD, BA, VAMG); and (iii) writing of the manuscript (AN, MS, BD, VAMG). All authors commented on the manuscript.

