## Supplementary Figures for "A new twist on bacterial motility – two distinct type IV pili revealed by cryoEM"

**Supplementary Information**

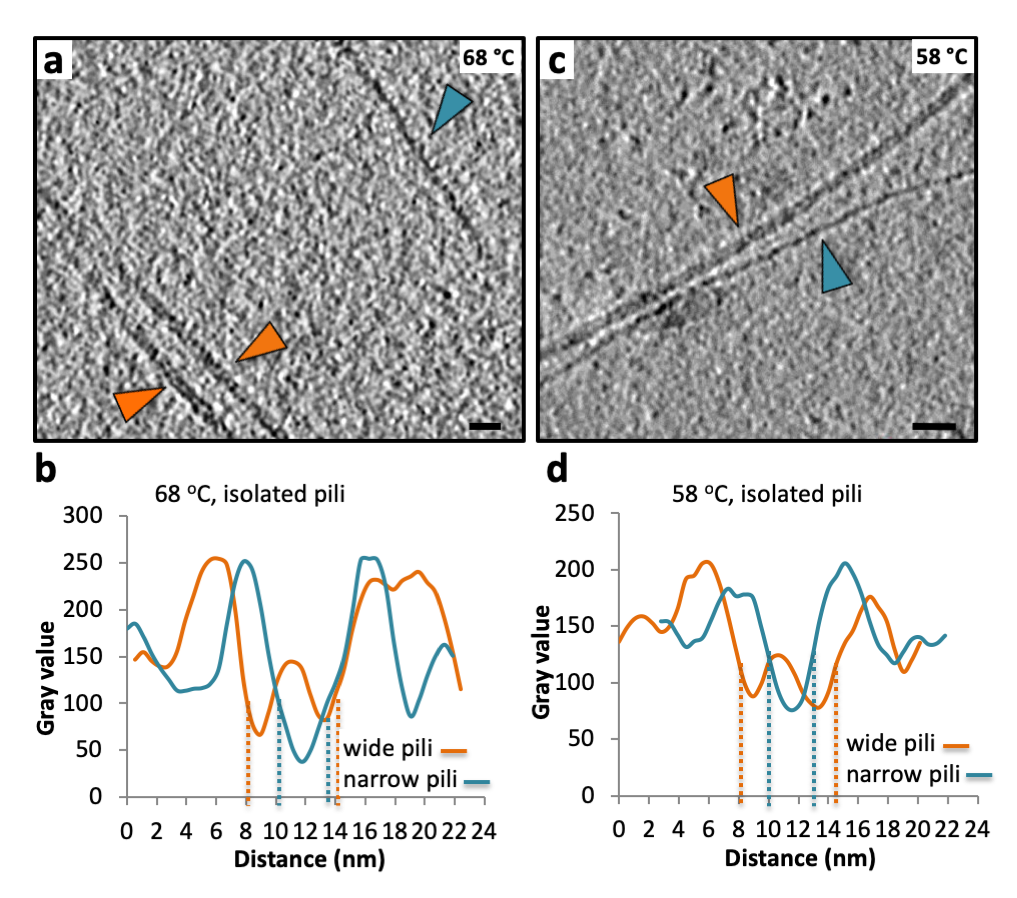

**Fig. S1: Purified pilus preparations contain both wide and narrow forms**

**a,** Tomographic slice through isolated pili from *T. thermophilus* grown at 68 °C shows both wide (orange arrowheads) and narrow pili (teal arrowhead). Scale bar, 20 nm.

**b,** Plot profile of grey values used to determine the diameter of two different pili, isolated from cells grown at 68 °C.

**c,** Tomographic slice through isolated pili from *T. thermophilus,* grown at 58 °C shows both wide (orange arrowhead) and narrow (teal arrowhead) pili. Scale bar, 20 nm.

**d,** Plot profile of grey values used to determine the diameter of two different pili, isolated from cells grown at 58 °C.

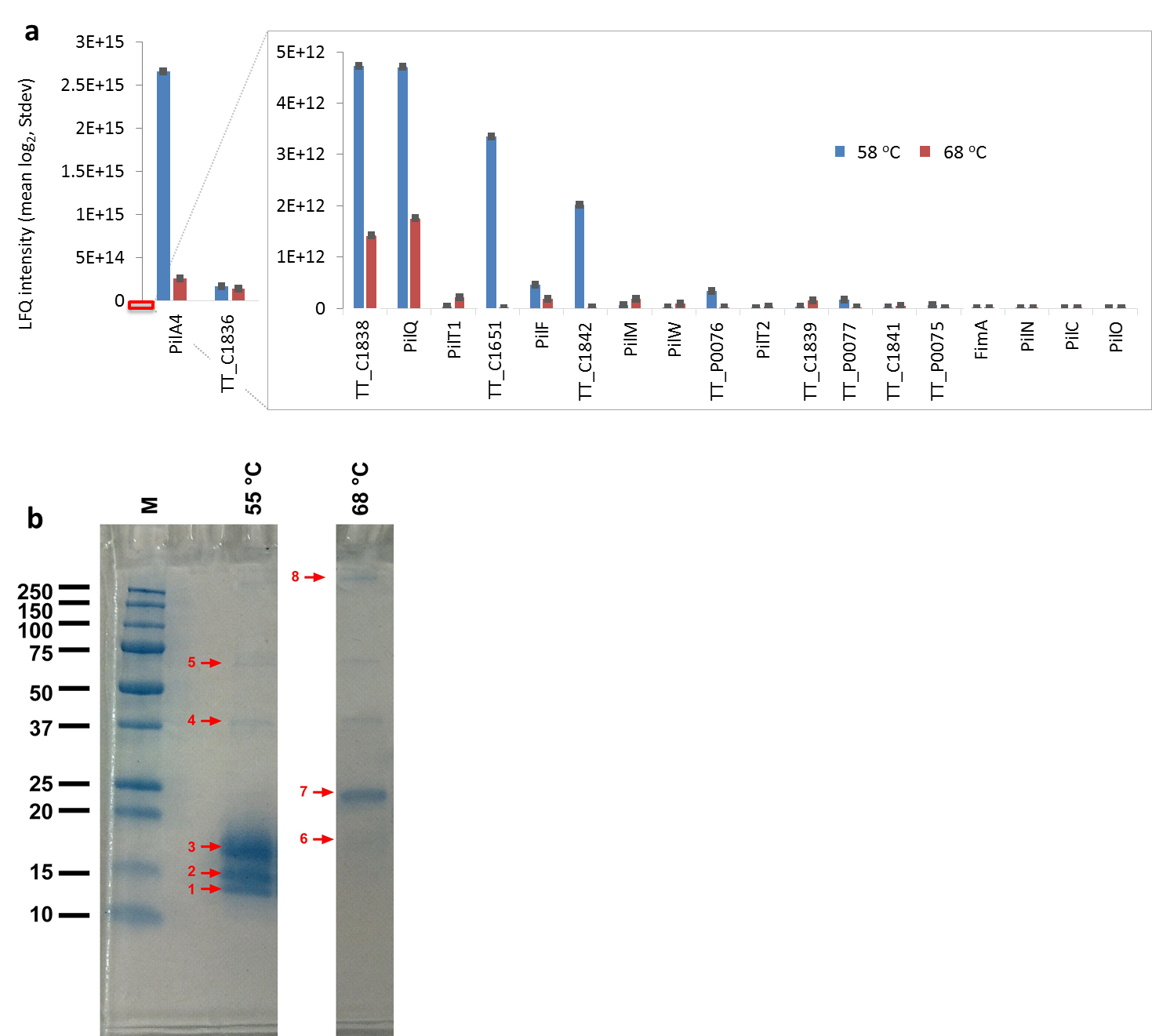

**Fig. S2: Mass spectrometry of isolated pili**

**a,** MS of isolated pili. LFQ intensity analysis showing the relative abundance of different pilin-like and pilus associated proteins at 58 °C and 68 °C.

**b,** Gel-based MS of isolated pili. Pili isolated from cells grown at 55 °C or 68 °C were purified and separated by SDS-PAGE. The numbered bands were analysed by MS and the most abundant proteins in each band are listed in Table S1.

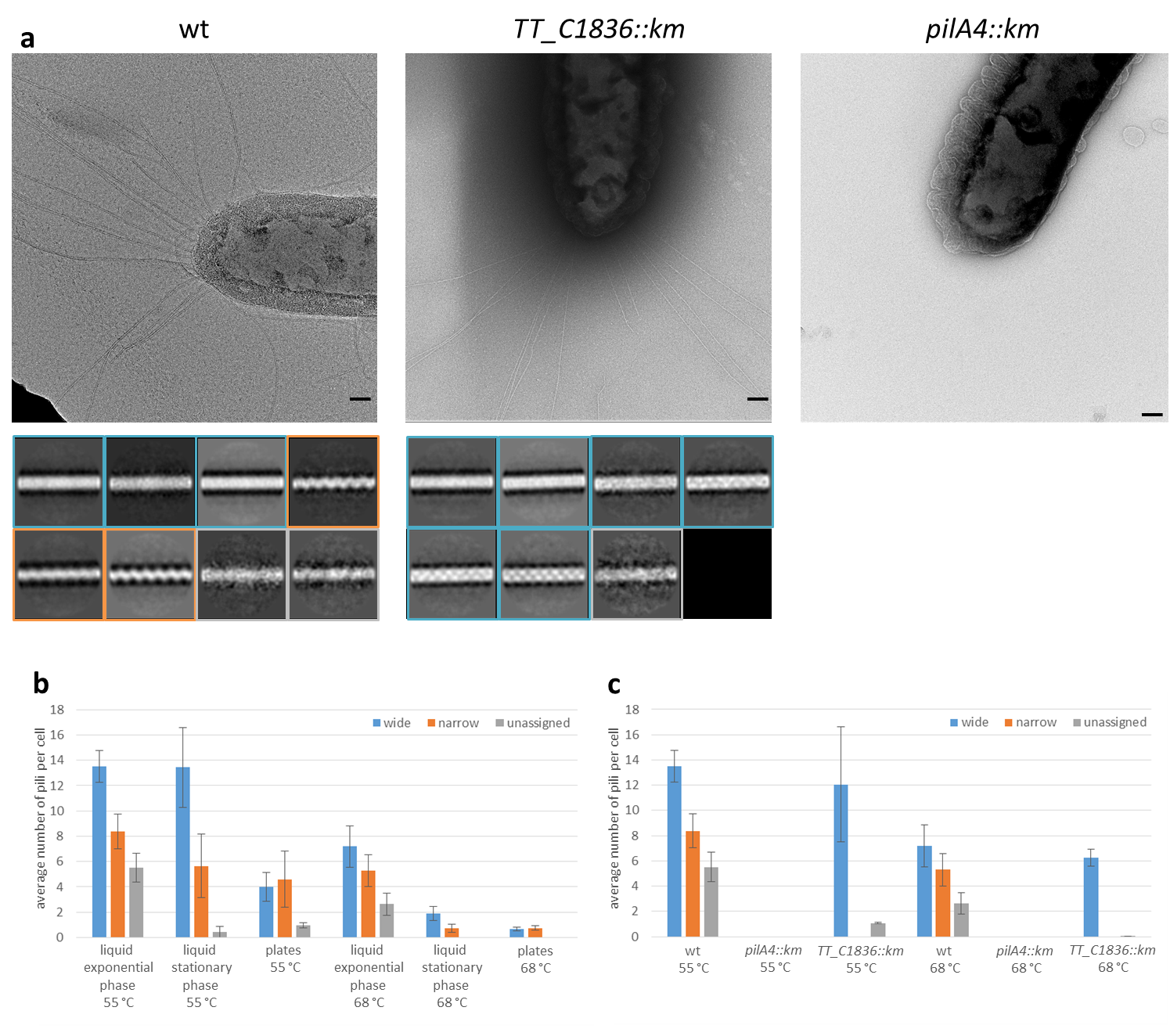

**Fig. S3: Analysis of the abundance of wide and narrow pili**

**a,** Top panels, representative electron micrographs of negatively stained *T. thermophilus* cells show long flexible filaments for wild type and *TT_C1836::km cells*. Scale bars, 100 nm. Bottom panels, examples of 2D class averages of pili from electron micrographs of whole cells. Class averages were assigned to wide pili (teal boxes) or narrow pili (orange boxes). Some filaments could not be assigned to classes reliably (grey boxes). The number of particles in each class was used for further evaluation in **b** and **c**. Box size, 35 nm.

**b,** Number and type of pilus assembled for wild-type cells under different growth conditions. Error bars represent the standard error from three independent experiments.

**c,** Number and type of pilus analysed for mutants grown to exponential phase. Error bars represent the standard error from three independent experiments.

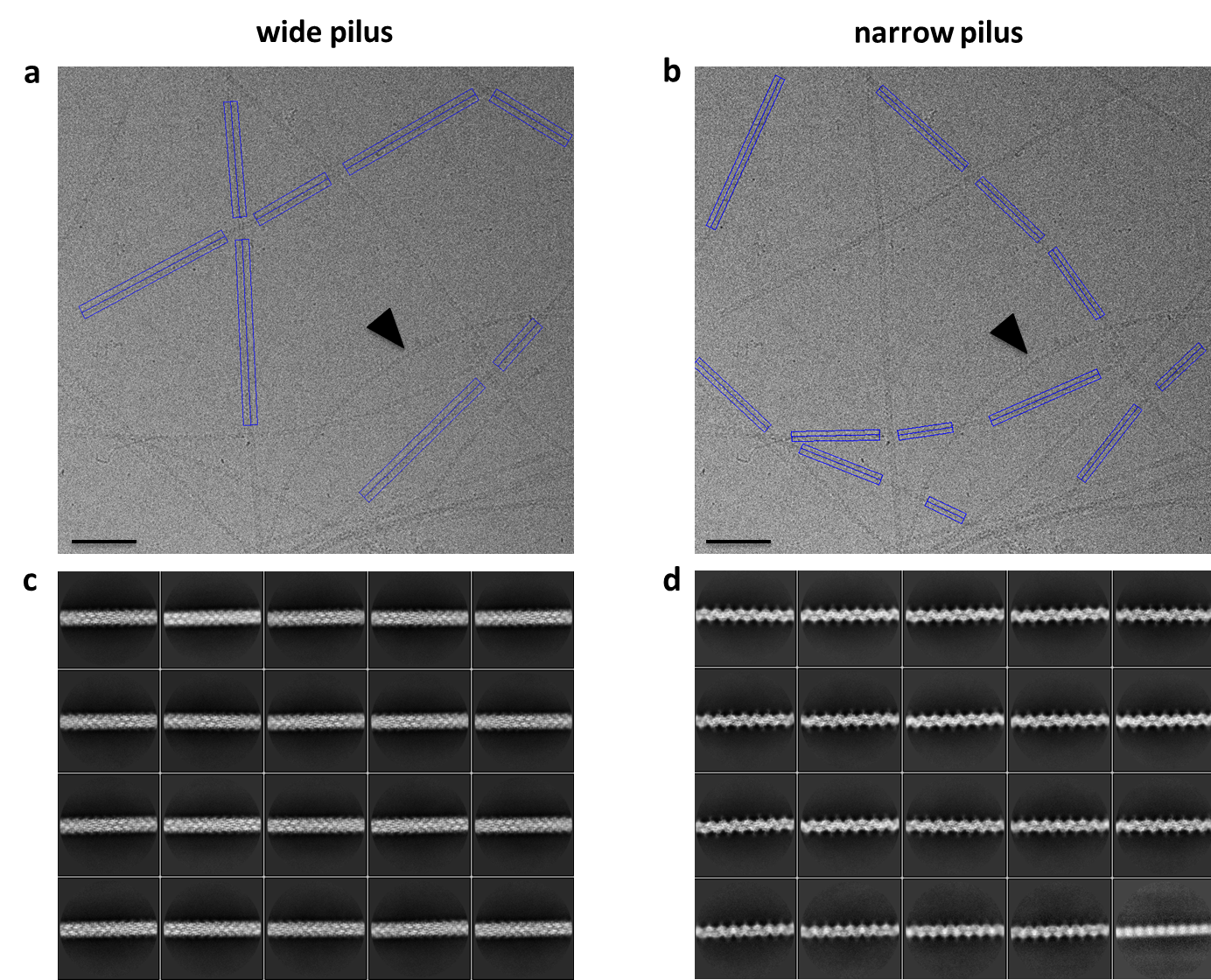

**Fig. S4: Raw CryoEM data and 2D classes**

**a, b,** CryoEM micrograph with picked wide (**a**) and narrow (**b**) pili. The arrowhead indicates a narrow pilus with a high degree of curvature that was often observed for this type of filament. Scale bars, 50 nm.

**c, d,** 2D classes of wide (**c**) and narrow (**d**) pili. Box size, 40 nm.

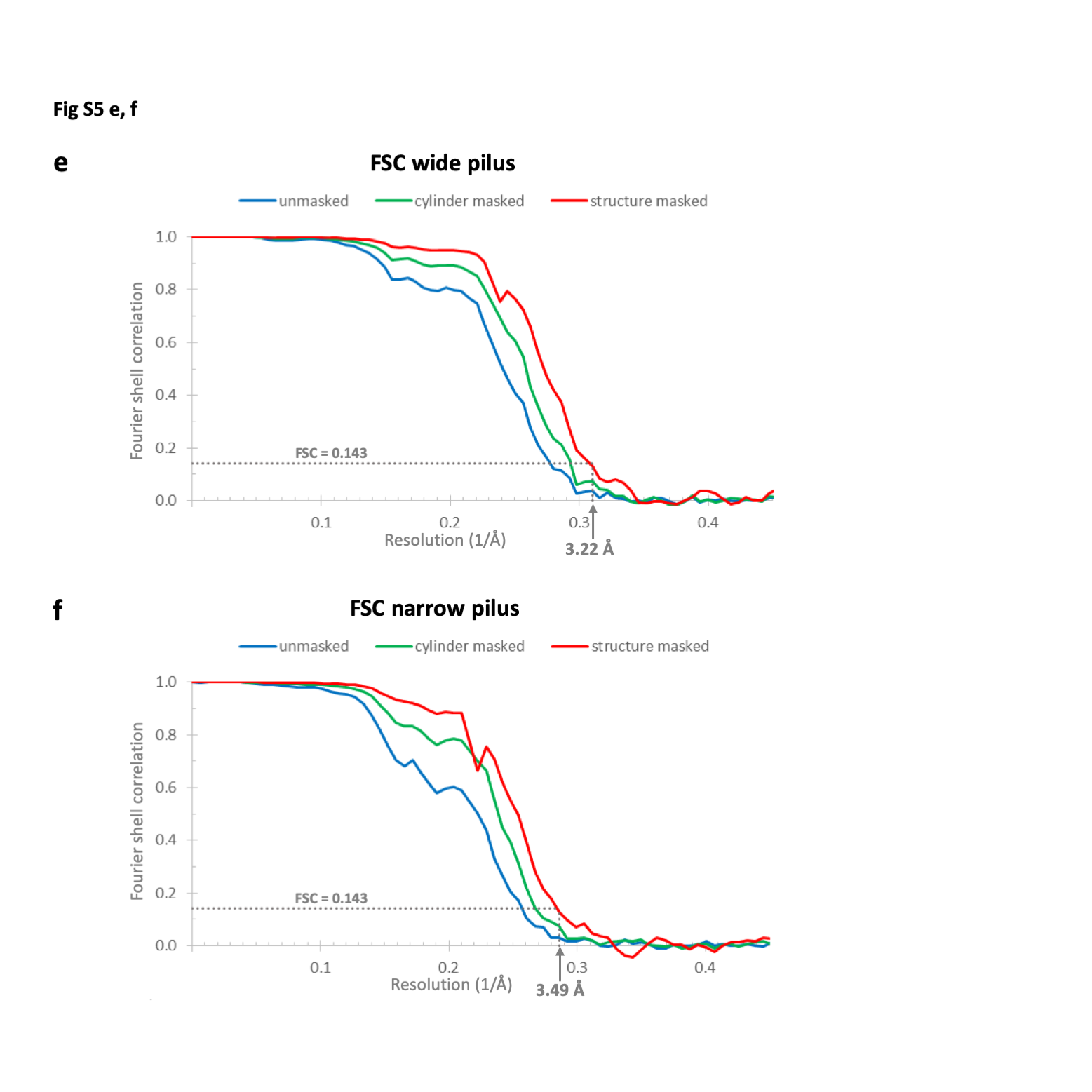

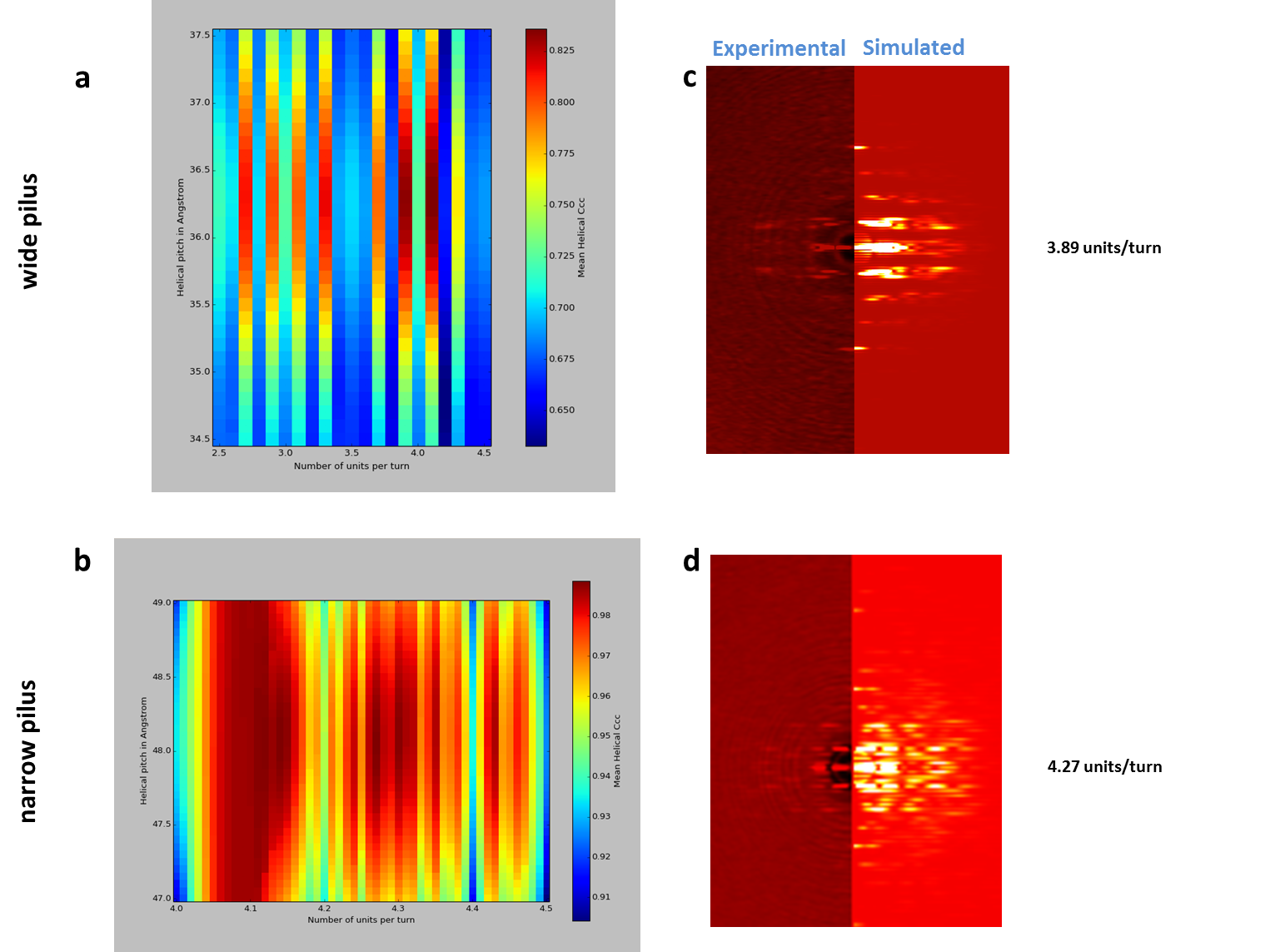
**Fig. S5: Helical symmetry determination using SPRING**

**a, b,** Segclassreconstruct (SPRING) results for wide (**a**) and narrow (**b**) pili. For wide pili a high correlation was observed for 4.1 and 3.89 subunits per turn and a helical pitch of ~36.3 Å (**a**). For narrow pili a high correlation was observed for 4.11, 4.14, 4.27 or 4.30 subunits per turn and a helical pitch of ~48.1 Å (**b**).

**c, d,** Agreement between layer line positions in experimental and simulated Fourier transforms for wide (**c**) and narrow (**d**) pili.

**e, f,** FSC curves for wide (**e**) and narrow (**f**) pili. The calculated final maps were determined at 3.22 Å resolution from 98,415 asymmetric units for wide filaments (**e**), and 3.49 Å from 76,866 asymmetric units for narrow filaments (**f**) using Fourier shell correlation (structure-masked, 0.143 cut-off). Blue curve, unmasked FSC; green curve, cylinder-masked FSC; red curve, structure-masked FSC.

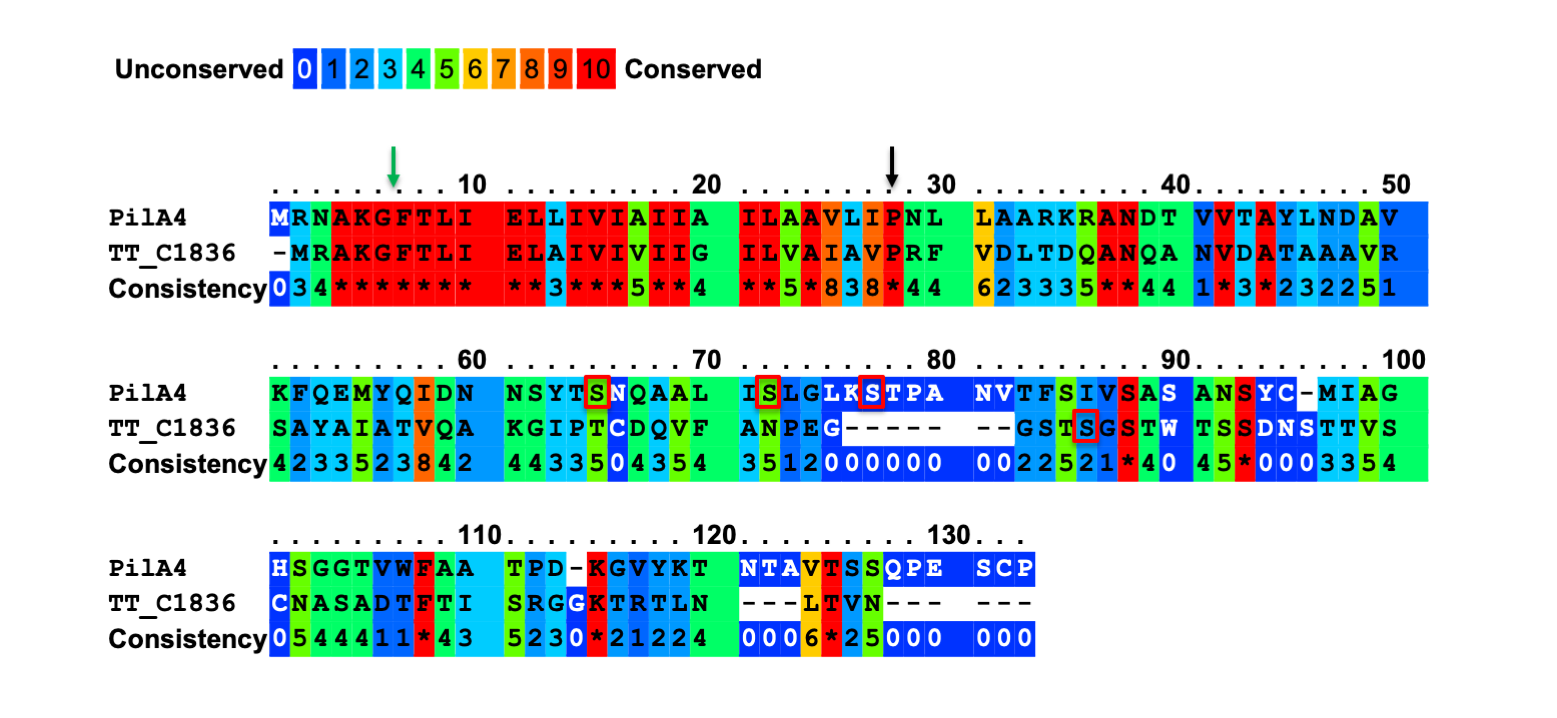

**Fig. S6: Sequence alignment between PilA4 and TT_C1836**
The cleavage site of prepilin peptidase (green arrow), the conserved Pro22 (black arrow) and putative glycosylation sites (red boxes) are shown.

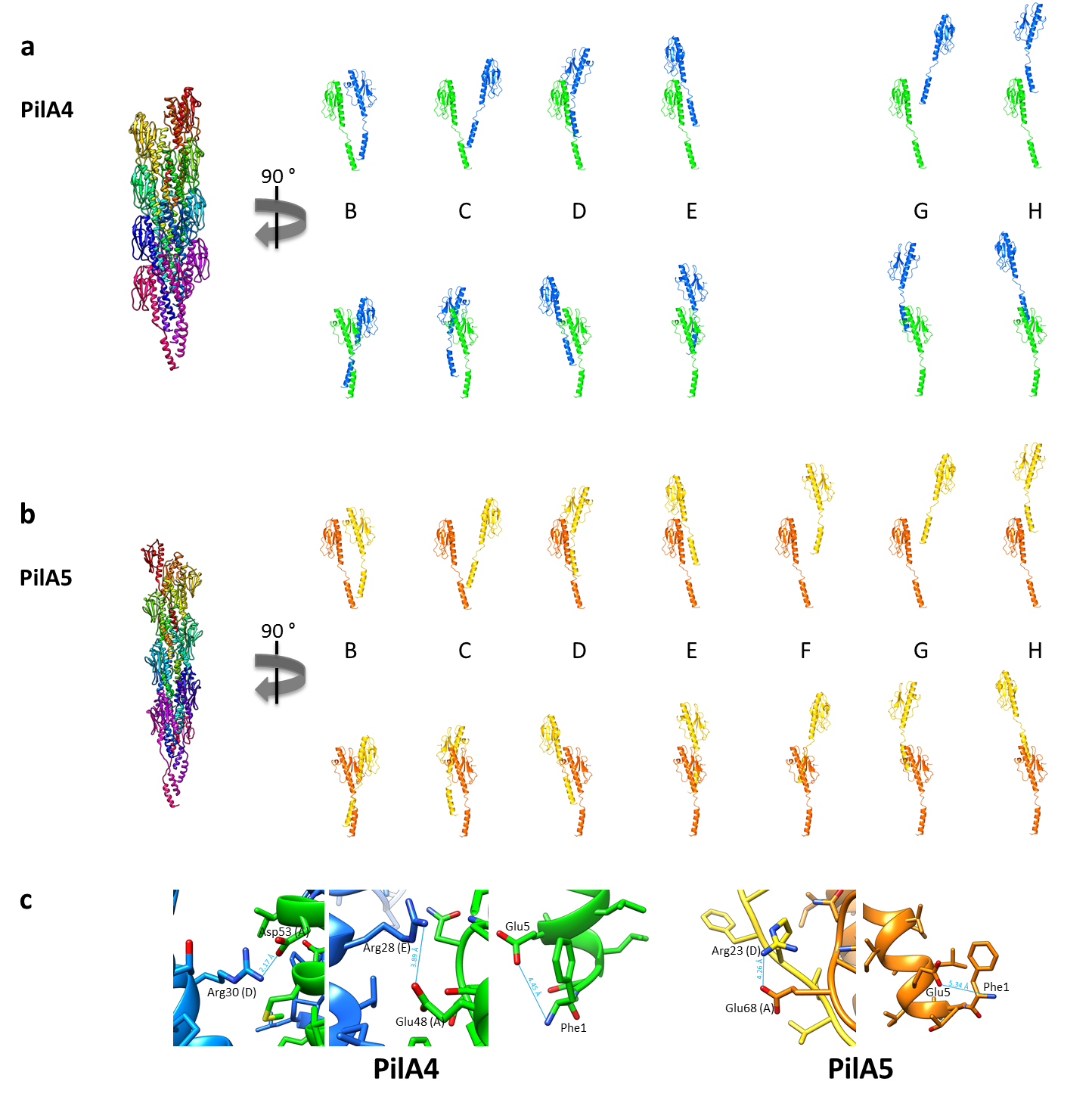

**Fig. S7: Intermolecular interactions within the filaments**

**a,** Ribbon representation of a 15mer wide filament (left) and of one subunit (subunit A, green) with another single subunit within the filament (blue, subunit B = +1 in the filament… subunit H = +7 in the filament).

**b,** Ribbon representation of a 15mer narrow filament (left) and one subunit (subunit A, orange) with another single subunit within the filament (yellow, subunit B = +1 in the filament… subunit H = +7 in the filament).

**c,** Close-ups of salt bridges within wide (left) and narrow (right) filaments. Same colour code as in (A, B).

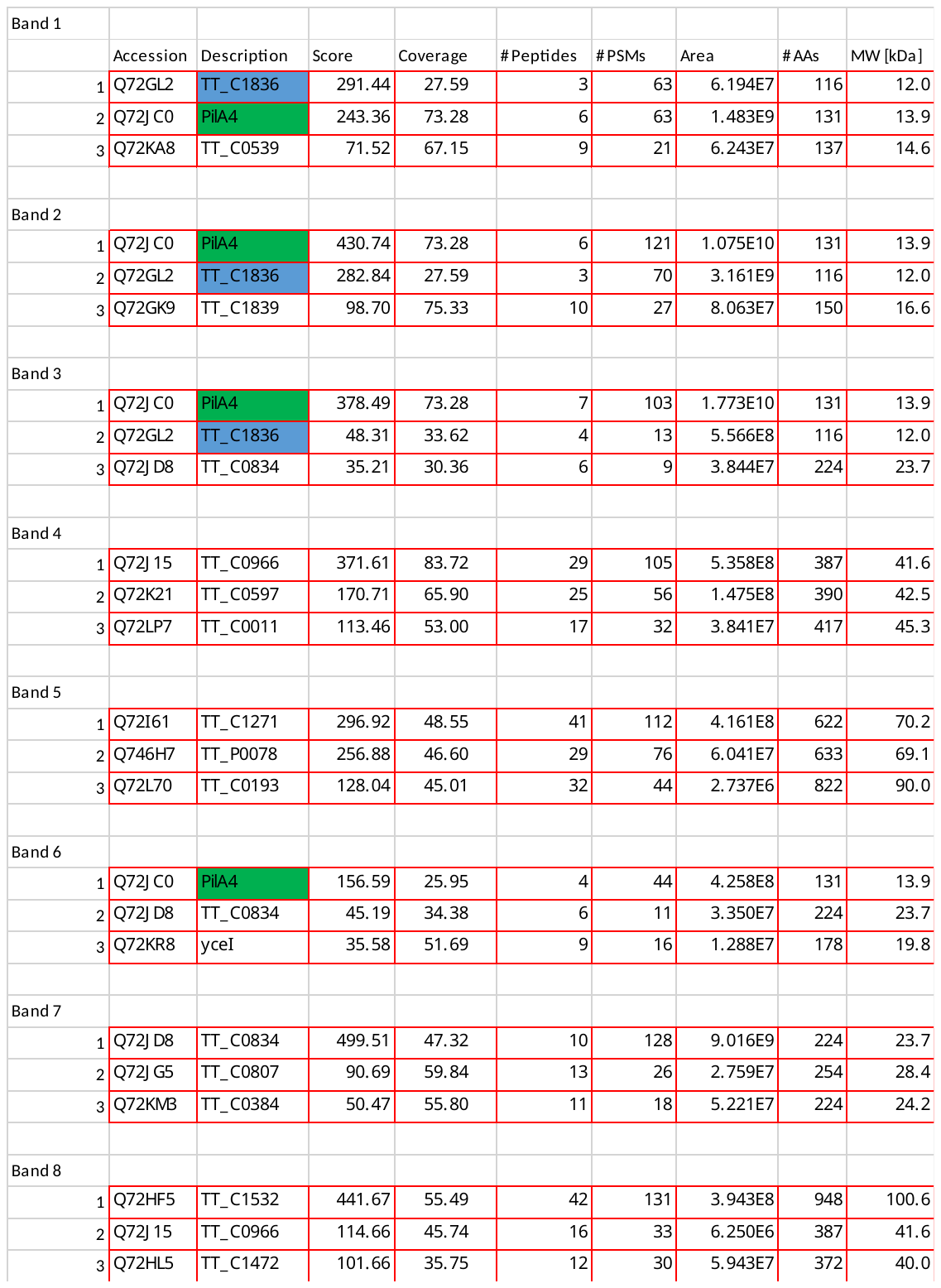

**Table S1: Proteomics of isolated pili samples (SDS-PAGE)**

Results of gel-based MS (Fig. S2b). For each band the proteins with the highest score were listed. PilA4 and TT_C1836 are highlighted in green and blue, respectively.
